# A Novel Design of Transcriptional Factor-mediated Dynamic Control of DNA Recombination

**DOI:** 10.1101/2020.10.20.346668

**Authors:** Jiayang Li, Yihao Zhang, Yeqing Zong

## Abstract

Genetic regulation is achieved by monitoring multiple levels of gene expression, from transcription to protein interaction. Unlike common temporary transcriptional regulation methods such as the use of inducible promoters, integrases permanently edit DNA sequences. Integrases, however, require especially strict regulation when implemented in synthetic genetic systems because of the irreversible result. Here we propose to improve the regulation of site-specific integrase-based genetic system by dynamically hiding one of the attB/P sites that are essential for recombination with transcriptional factors. After effectively suppressing excessive recombination, we also validated the necessity of each of the essential components in our transcriptional factor-controlled recombination (TFCR) system. The system applied transcriptional-level regulators directly on controlling the activity of existing non-transcriptional-level trans-factor proteins by inhibiting its binding to the cis-regulatory elements. We anticipate my results to provide greater robustness for integrase components to enable safer use in systems as well as be a starting point for future cross-expression level gene regulations.

## Introduction

Found in bacteriophages, integrases, a kind of site-specific recombinases, naturally function to insert phage DNA into bacteria chromosome. Specifically recognizing two specific sites (attB and attP), integrase recombines the DNA between them. Capable of permanently excising or inversing DNA regions, integrases have been applied in diverse biological systems. In classical molecular operations, integrases were used to perform site-specific in vivo assembly of genes (Colloms, Merrick et al. 2013). Researchers even showed the possibility to target sites in human cells with phage integrases and recombine them (Groth, Olivares et al. 2000). In the field of synthetic biology, its ability to irreversibly change the genetic circuit enabled researchers to permanently record events and the order in which they occur in a cell (Roquet, Soleimany et al. 2016). It has been found that some integrases also possess high orthogonality, enabling them to be used together within a single complex system as biological registers to record over digital information and to create state machines in cells (Yang, Nielsen et al. 2014). Other researches have also shown that the integrase system can be used as a rewritable addressable data module to store digital information within chromosomes (Bonnet, Subsoontorn et al. 2012).

While utilizing integrase’s unique characteristics, a problem that cannot be ignored is the excessive expression of integrase. Without proper control, recombination can irreversibly happen without manual induction in artificial genetic circuits, which reduces the controllability of the system and weakens precise regulation.

The hyperactivity of integrases can be suppressed by adjusting components at different levels of gene expression: transcription, translation, and protein interaction. Previous systems implementing integrases have applied fine regulation on the component through methods such as using tightly regulated promoters like pBAD (Roquet, Soleimany et al. 2016), decreasing the integrase translation by switching start codon5, or implementing a riboregulator (Siuti, Yazbek et al. 2013).

Here, we provide a design to effectively regulate recombination on a different level by introducing transcriptional factors with the ability of high-order oligomerization. According to the common model, recombination does not occur without recognition of both recognition sites by integrases. Taking advantage of this theory, we attempted to disable interactions between an integrase and its attB site. Transcriptional factors such as repressors naturally have the ability to disable interactions between RNA polymerase and promoter region; taking advantage of the DNA looping ability of transcriptional factors with oligomerize domain, we attempted to insulate attB sites with the loop. We also established a biophysical model to mimic the mechanism of the competitive inhibitor transcriptional factors (TFs) and proposed several pathways for further optimization through sensitivity analysis.

## Materials & Methods

### Construction of integrase plasmids and transcriptional factor plasmids

Plasmids used in this study were constructed using methods including restriction enzyme cloning, Golden Gate assembly, or Gibson assembly. Correct colonies are selected using antibiotic selection and examined via DNA sequencing.

### Construction of reporter plasmids

The vector for the reporter plasmids was assembled from a BAC vector derived from pcc1BAC. Superfold-GFP expression unit from the pPT plasmid (Zong, Zhang et al. 2017) was added onto the vector through BioBrick Standard Assembly. The region with T7 promoter and φC31 attP site was added by PCR with primers carrying over 20 bp homologous arm to the vector as well as 20-30 bp overlapping the new region. The plasmid was transformed into competent *Escherichia. coli*, verified through sequencing, and then extracted and purified. The region with the operators (or meaningless sequence with the same length) and the attB site was constructed from PCR with primers and PCR Phusion. PCR was carried with two primers with over 20 bp overlapping region without a plasmid template. The product of this PCR was then used in another PCR, amplified by the old reverse primer and a new forward primer with 20 bp overlapping region with the old forward primer to elongate the length of the previous PCR product. The final PCR product was designed to carry two restriction enzyme recognition sites with three protection base pairs at each of the two ends. These two restriction enzyme recognition sites enabled them to be inserted into the formerly constructed reporter plasmid at the appropriate location.

All plasmid maps are provided in supplementary material.

### Strain preparation

Reporter plasmids (pRR), TF plasmids (pTF) and recombinase plasmids (pR) were constructed and extracted separately. For each combination, a reporter plasmid was transformed into *E. coli* DH10B in which a constitutively expressing T7 RNA polymerase module is integrated into the genome (Zong, Zhang et al. 2017) first. The strains were then made competent and the TF plasmid and recombinase (integrase) plasmid were co-transformed into these competent strains carrying reporter plasmids (Figure 1). The final *E. coli* strains with three plasmids were grown overnight on LB agar plates in 37 °C with antibiotic selection of kanamycin, ampicillin, and chloramphenicol. The expression of integrase and TF could be controlled by addition of inducers (cumate and IPTG).

**Figure 1.**
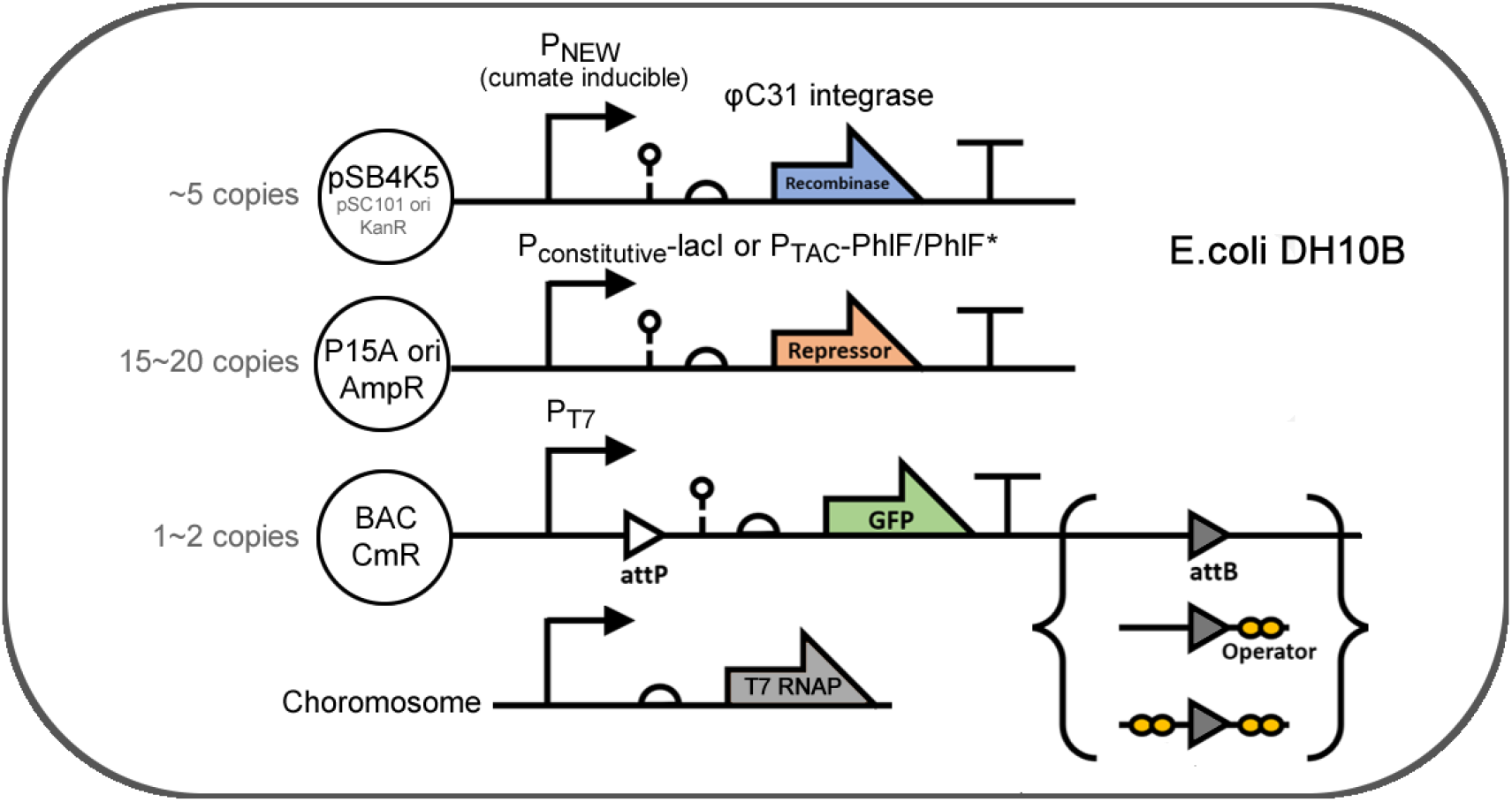
Plasmids and strain used in this research

### Fluorescence measurement

Regions or colonies of *E. coli* were selected from the agar plate and inoculated into liquid M9 medium with antibiotic selection in 37 °C as well.

Fluorescence of groups in the experiments was both observed with the naked eye or fluorescent microscope and quantitatively measured through flow cytometry.

For flow cytometry measurement, the detailed procedures are as follows:

1. Co-transfer the integrase-expressing plasmid, TF plasmid, and reporter plasmid into the host *E. coli* strain.
2. Inoculate monoclonal colony in the M9 supplemented medium.
3. The cell cultures were diluted 1000-fold with M9 supplemented medium of various inducer (IPTG and/or cumate, depending on the experiment) concentrations, and were incubated for 20h.
4. 3μl samples of each culture were transferred to a new plate containing 200 μl per well of PBS supplemented with 2 mg/μL kanamycin to terminate protein expression. The fluorescence distribution of each sample was assayed using a flow cytometer with appropriate voltage settings. The ratio of fluorescent cells and the arithmetical mean of each sample were determined using FlowJo software.

### Media and buffer

LB medium: 10 g/L tryptone, 5 g/L yeast extract, and 10 g/L NaCl. For agar plates, 15 g/L agar was added.

M9 supplemented medium: 6.8 g/L, Na_2_HPO_4_, 3 g/L KH_2_PO_4_, 0.5 g/L NaCl, 1 g/L NH_4_Cl, 0.34 g/L thiamine, 0.2% casamino acids, 0.4% glucose, 2 mM MgSO_4_, and 100 μM CaCl_2_.

Antibiotic concentrations: the media contained ampicillin at a final concentration of 100 μg/mL, chloramphenicol at a final concentration of 25 μg/mL, kanamycin-sulfate at a final concentration of 50 μg/mL. The indicated concentrations of antibiotics were used for both LB and M9 media.Inducers: isopropyl-d-1-thiogalactopyranoside (IPTG) at a final concentration of 500 Μm. A 4-isopropylbenzoate (cumate) gradient was prepared at 0, 10, 50, 100, 200, and 400 μM final concentrations.

PBS Buffer: 8 g/L NaCl, 0.2 g/L KCl, 1.44 g/L Na_2_HPO_4_, and 0.24 g/L KH_2_PO_4_. Kanamycin (2 mg/mL) was added to the PBS before sampling in order to terminate protein expression.

## Results

### The strategies of system design and measurment

The transcriptional factor-controlled recombination (TFCR) system we designed contains three plasmids: a recombinase plasmid that expresses integrase φC31, a TF plasmid that results in induced or constitutive TF protein expression, and a reporter plasmid that indicates recombinant reaction quantitatively.

The basic structure of the reporter plasmid consists of a T7 promoter, an attP site, GFP, a terminator and an attB site parallel to the attP site. We attempted to avoid integrase activity by putting operator sites adjoined to the attB site which is necessary for recombination according to common models (Smith, Brown et al. 2010, Bonnet, Subsoontorn et al. 2012). Three types of reporter plasmids are set up, each containing none, one or two operator sites around the attB site. For the groups with two operators, one operator is placed on each side of the attB site. The spacing between two operators was set around a multiple of 10 bp to make sure that both operators were located on the same side of the DNA double helix. For the groups with zero or one operator sites, meaningless sequences with the length of the operators are added at the location of the operators to maintain equal distances between the attP and attB sites in each group. We assume that when the TFs bind to the operator sites around the attB region, the integrase becomes no longer able to recognize the attP sites, which prevents further recombination.

Without the interference of TFs, the two parallel recombination sites are exposed to integrases to enable recombination, resulting in excision of the region of DNA between the two sites containing the green fluorescent protein expressing module. This would result in a decreasing proportion of high-fluorescence *E. coli* in the population. Therefore, in flow cytometry measurement, a lower percentage of cells at high-fluorescence is considered as the identification of high integrase activity, and vice versa (Figure 2).

**Figure 2.**
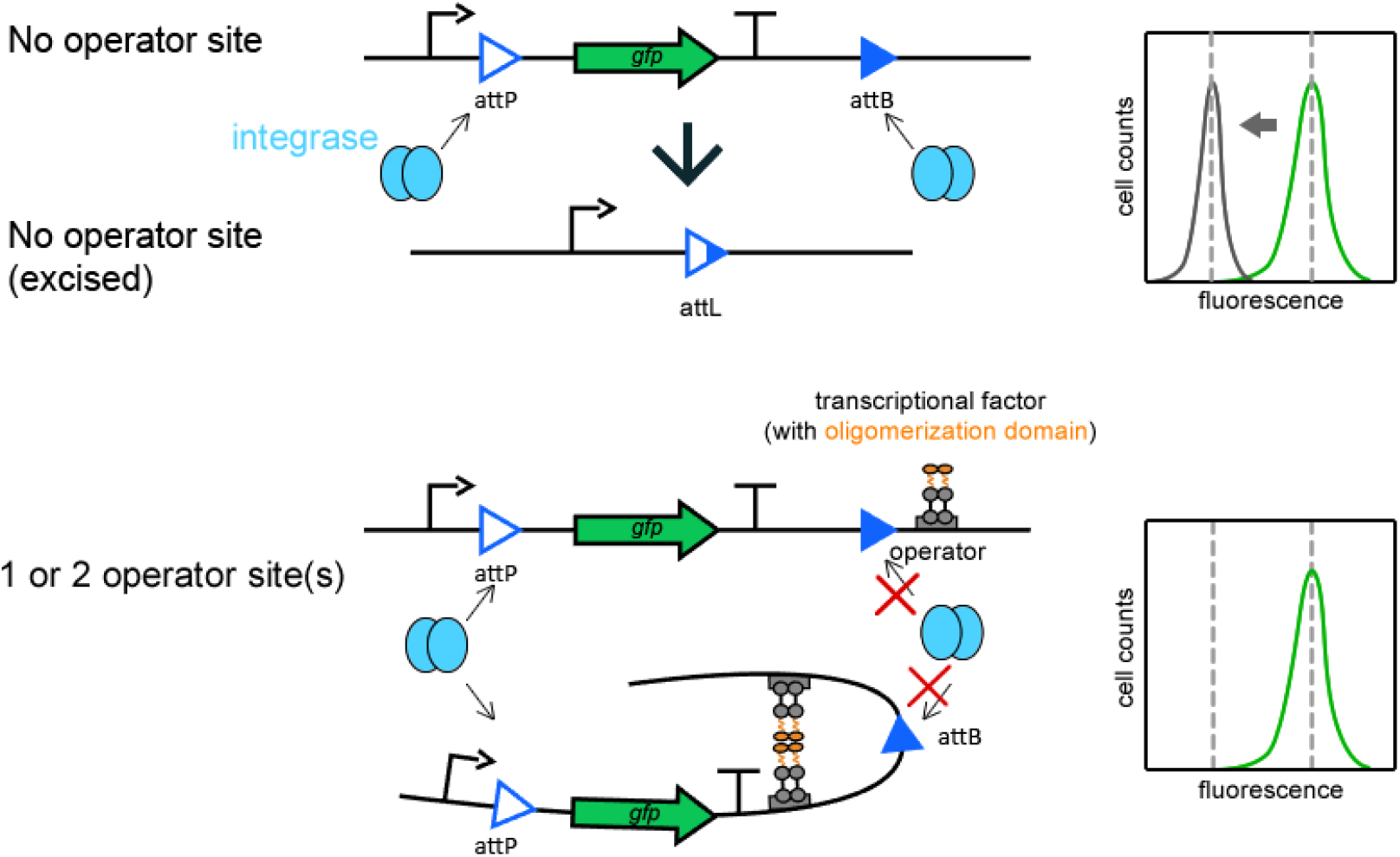
The design and mechanism of TFCR system. In the reporter plasmid with no operator sites, integrase can recognize the attP and attB sites and recombine them, excising the DNA region between the two sites. The circuit after excision is shown as “No operator site (excised)”. The diagram for 1 or 2 operator site(s) illustrates the proposed model of TF(s) with oligomerization domains in which TF(s) “blocked” the region of attB, disabling recognition and excision by integrases. Diagrams on the right represent expected fluorescent cell distribution of the systems on the left measured with flow cytometry assay.

**Figure 3.**
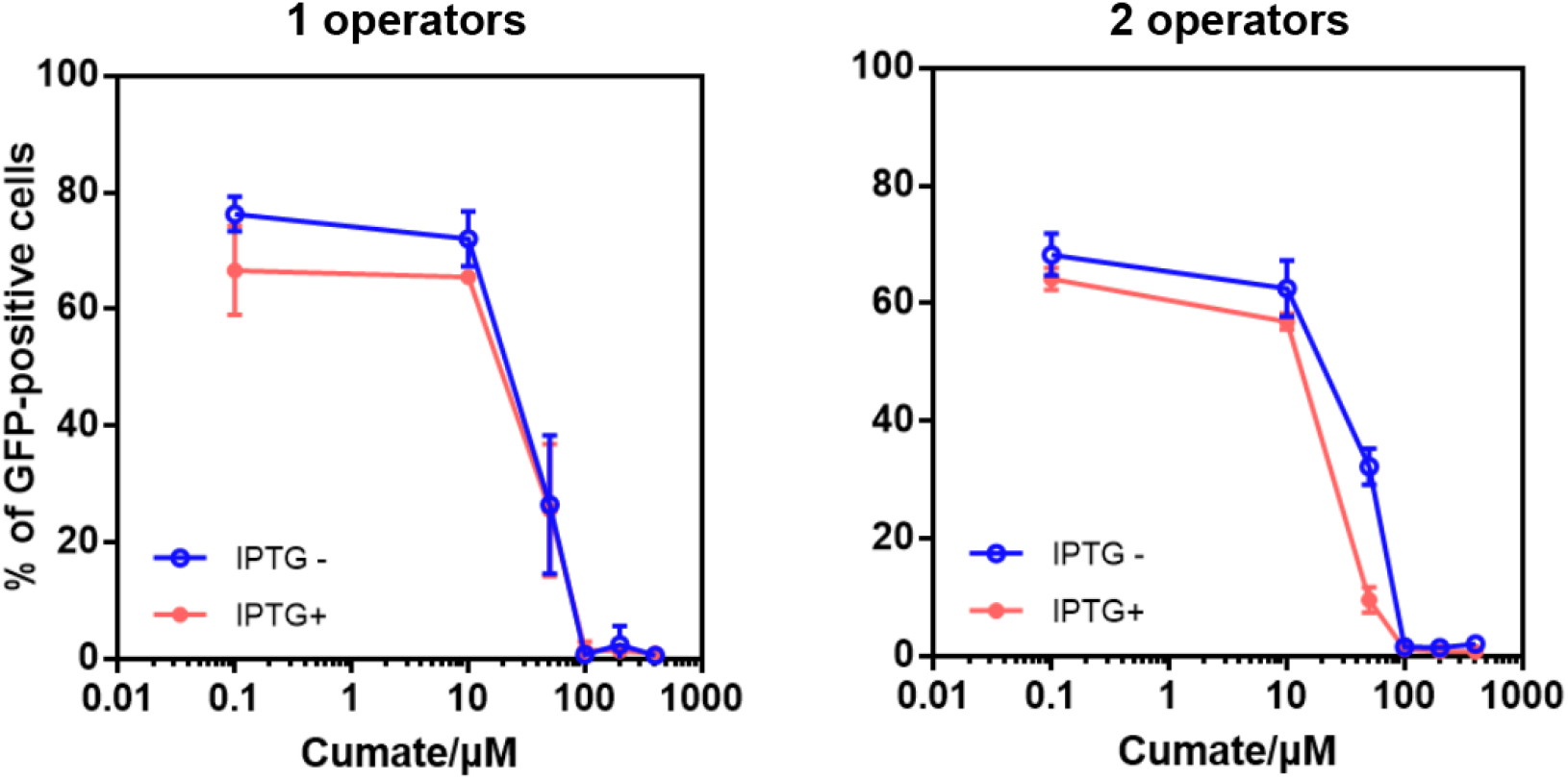
Testing the effect of the number of operators on the reduction of recombination. Two systems are set up, one with one lacO and the other with 2 lacO, to test the plausibility of repressing recombination with TFs. Recombination levels were measured at 0, 10, 50, 100, 200, 400 μM cuamte induction levels of the integrase φC31. Data represent the means ± SD from at least three replicate experiments conducted on different days.

### Proof of concept by testing the lacI-based TFCR system

According to the hypothesized DNA bending mechanism of these TFs, the strong blocking function requires both the operator sites to bind with each other and hide the att site in between. To test the plausibility of the TFCR system, we compared the performance of systems with one or two operator sites around the attB site. We chose the widely applied lacI TF and φC31 integrase for the experiment to test the effect of the number of operators on reduction of recombination rate. Due to the uncertain effect on recombination rate by the difference on the DNA sequence of trans-regulatory element, we focused on comparing results from groups with and without activated TF at different integrase expression levels consisted of identical systems. IPTG was used to inactivate the LacI protein.

The group with one operator site demonstrated little difference on recombination rate between groups with and without TF plasmid induction at high integrase induction levels, showing minimal repression activity by the LacI TF. The group with two operators IPTG-showed over 20 % reduction of recombination level at 50 μM cumate than the group with 2 operators IPTG+, while the group with 1 operators showed little difference. This demonstrated the plausibility of reducing integrase activity with the TFCR system.

Since the design with two operators demonstrated significantly better functions at than the design with one operator, we continued to use the two-operator set in the following experiments.

### Improving the TFCR system using high-performance TF

Despite the considerable difference was shown in the LacI-based experiment, it is hard to say that’s an “effective regulation”. We suspected that this may be due to the relatively weak repression ability of LacI. Thus, we replaced LacI with an artificial transcriptional factor PhlF* which was derived from PhlF and fused with an oligomerization domain from another TF protein CI434 to increase its binding performance (Hou, Zeng et al. 2018). The new TF plasmid harboring the PhlF* gene also implements an inducible P_TAC_ promoter to enable convenient control of TF expression. With PhlF*, two groups of experiments were set up to validate the necessity of each component in the improved TFCR system.

I tested the effect of PhlF* within the system. The experimental group consists of a reporter plasmid with two operators, the TF plasmid and the integrase plasmid. This group is treated with 0.5 mM IPTG to initiate the expression of the PhlF*. The second group harbors the same plasmids but is not induced with IPTG. Results from flow cytometry showed that at a representative high integrase induction level, the group with sufficient PhlF* expression (IPTG added) has little decrease in the percentage of high fluorescence cells compared to the group without sufficient PhlF* expression (no IPTG) (Figure 4A). It is reasonable to conclude that PhlF* played the important role in the inhibition of recombination in the system.

**Figure 4.**
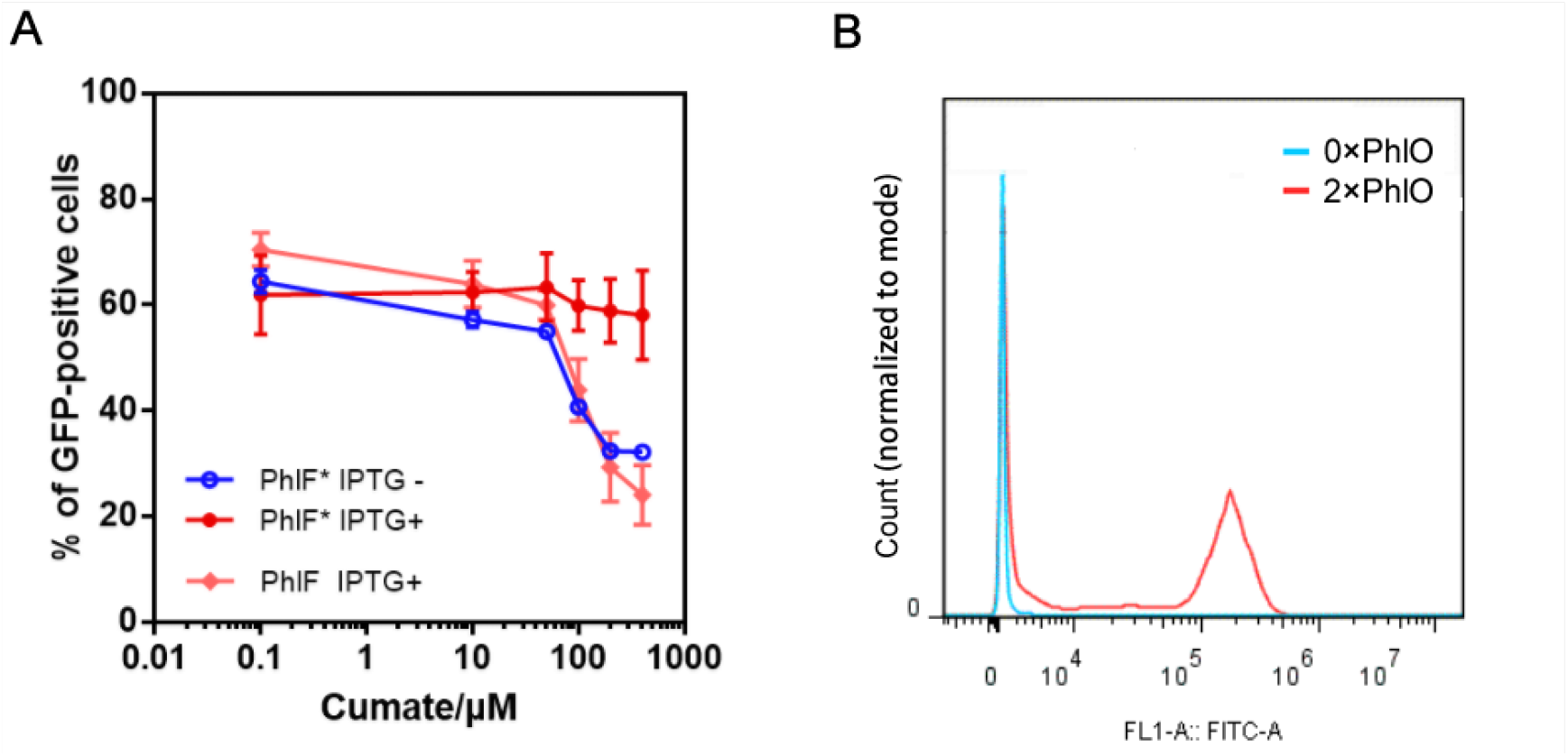
The PhlF*/PhlF-based TFCR system. A. Comparing results from systems with induced PhlF* (red), uninduced PhlF* (pink) and induced wild type PhlF (blue). Data represent the means ± SD from at least three replicate experiments conducted on different days. B. The representative fluorescent cell distributions of the groups with (red) or without (blue) PhlO. The IPTG concentrations were 0.5 mM, and the cumate concentrations were 400 μM.

In order to eliminate the possibility that the decrease of recombination is the result of other effects such as cytotoxicity caused by PhlF*, the necessity of PhlO, the operator of PhlF in the system was also verified. Two groups with all three plasmids are constructed, one with two operator sites on the reporter plasmid and the other with none. At the high expression level of integrase, and with IPTG induction, the group with no PhlO showed nearly zero percentage of high-fluorescence cells, suggesting the full activity of integrase (Figure 4B). Thus, we confirmed that the decrease in recombination is caused by the binding of PhlF* to PhlO.

In addition, we validated suspect of the artificially appended oligomerization domain as an essential requirement for the blocking of the recombinant site. A group with induced wild type PhlF replacing the engineered PhlF* in an otherwise identical system was set up and compared with the group with the PhlF* system. The group with the wild type PhlF demonstrated significantly higher recombination rates that were similar to those from the uninduced PhlF* system (Figure 4A). The results here matched the model of TFs with oligomerization domains and these domains are shown indispensable for proper function of the system.

## Modeling

In order to direct further optimization of the TFCR system, we established a biophysical model to mimic it. The core principle to the recombination-control of this system is based on the competitive inhibition mechanism of the TFs on the specific integrase recognition sites. This simplified process can be described with reference to the thermodynamic model for transcriptional regulation system (Zong, Zhang et al. 2017).

In thermodynamic equilibrium state, the probability for integrases to bind with an attP site (*p*_P_) can be expressed as a Hill function:

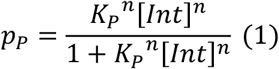

where *K*_*P*_, [*Int*], and *n* represent the binding constant between the integrase and the attP site, the intracellular concentration of integrases, and the Hill coefficient of the binding between integrases and attP site.

Meanwhile, the probability of binding between the integrase and the attB site *p*_B_ can be expressed as:

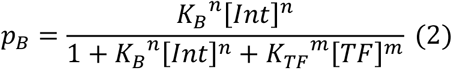

where *K*_*P*_, *K*_*TF*_ and m represent the binding constant between the integrase and the attP site, the binding constant between the TF and the attP site and the Hill coefficient of the binding between TF(s) and attB-Operator(s) sites.

With the assumption that the DNA recombination rate *r* positively correlates to the probability of the two events described above, with the ratio α, we can represent the DNA recombination rate as:

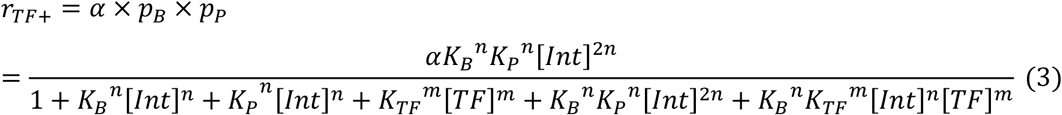

While without the presence of TFs in the system, the DNA recombination rate can be expressed as:

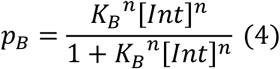

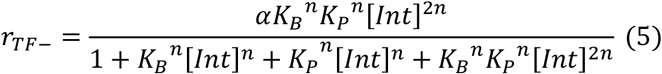

With all other conditions constant, it is obvious that *r*_*TF*−_ < *r*_*TF*+_.

In order to maximize the control on recombination rate, we need *r*_*TF*+_ << *r*_*TF*−_. This requires 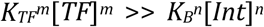, with maximal binding of TFs to operators and minimal binding of integrases to attB sites. This can be done through adjusting the two components.

The values of [*TF*], *m*, and *K*_*TF*_ can be increased. The concentration of TFs in cells can be increased through adjusting expression levels until they create excess burden for the cell and become cytotoxic. With this limitation, the increase of *K*_*TF*_ and *m*, inherent characteristics of the TFs become more significance. Therefore, choosing TFs with high-order oligomerization domains would be optimal. Experimental data above had shown that the TF PhlF* with higher-order oligomerization abilities resulted in more tightly regulation of integrases than LacI and PhlF with lower-order or none oligomerization domains. More TF parts with oligomerization domains are available to provide variability on functions of practical system depending on the desired level of expression (Hou, Zeng et al. 2018). Increasing value *m* through increasing the number of operators may also be plausible.

On the other hand, *K*_*P*_, [*Int*], and *n*, variables related to integrase functions can be adjusted. As a proof, we had replaced the P_NEW_ promoter which is upstream to the integrase gene with a strong constitutive promoter, resulting in high recombination rate even with the high expression level of PhlF*(Data not shown). Likewise, adjusting them can also decrease recombination activity. Decreasing integrase concentration and fine-tuning the attB sequence to get a lower affinity to integrases are currently available regulation methods.

Regulations on different levels multiply the possibility of achieving an ideal system with a finely adjusted signal processing function. Therefore, in a practical scenario, the TFCR system performs best supplemented by other typical strategies.

## Discussion

In former studies, integrase activity was controlled at transcriptional and translational levels with methods such as changing promoter strength, finetuning RBS, and using tightly regulated promoters (Roquet, Soleimany et al. 2016, Bonnet, Subsoontorn et al. 2012). In this project, we suggested the blocking of recognition site with operators as a new level of integrase regulation. This level is orthogonal to the other regulation methods, adding a new possible approach to be used with the above-mentioned methods in the same system. With the addition of each level of regulation, the efficiency of regulation multiplies, reducing the possibility and extent of expression leakage. Therefore, through increasing robustness and controllability of integrases, this project offers them greater potential to be applied for gene regulation in engineered biological systems.

Our design enables synthetic biological systems to apply controllable removal of DNA sequences. In a patented biological manufacturing system, integrases can be programmed to delete the patented gene components when the cells leave the desired conditions and prevent the theft of intellectual properties. A similar system can be applied to pollutive biological manufacturing factories; with gene contents which express pollutive chemicals removed by integrases when the cells leave ideal incubation conditions, biochemical pollution to natural water sources can be reduced efficiently.

On the other hand, we explored more potentials of TFs, which are commonly used only for the regulation of promoters, operators, and repressors that control downstream transcription. Formerly used only for direct regulation of transcription, TFs were now found the possibility to regulate trans-factor protein activity by interacting with the DNA region containing cis-regulatory elements. Its function of regulating integrase activity demonstrates its potential of interacting with other cis-regulatory elements to enable the locking of DNA regions to prevent unwanted reactions with substances in the environment, allowing further complex signal processing through the addition of a new controllable component.

Apart from this, the results of this project were consistent with the model established by previous researches on integrase (Smith, Brown et al. 2010, Bonnet, Subsoontorn et al. 2012). According to the model of recombination by large serine recombinases, two integrase dimers each need to bind with an attB/P site to perform recombination. In the experiments of our project, recombination ceased after disabling the recognition of one of the two sites. This result satisfies the former established model by verifying the necessity of the integrase and both attB/P sites in recombination.

Furthermore, we took advantage of independent parts artificially modified by synthetic biologists. Replacing the natural LacI repressor with the engineered PhlF* repressors with oligomerization domains, we achieved a more rigorous repression activity. Utilizing artificially modified or assembled vectors with standard BioBrick components, we conveniently constructed our plasmids with appropriate antibiotic resistance and number of copies. Building off the database with works by former researchers, we added the new integrase system to provide another option for signal processing for future works.

