## Supplementary material for "A Novel Design of Transcriptional Factor-mediated Dynamic Control of DNA Recombination": Supp_A_Novel_Design_of_Transcriptional_Factor-mediated Dynamic_Control of_DNA_Recombination.pdf

### Plasmid Maps

All the plasmid map images were generated by SnapGene®.

**Recombinase plasmids**

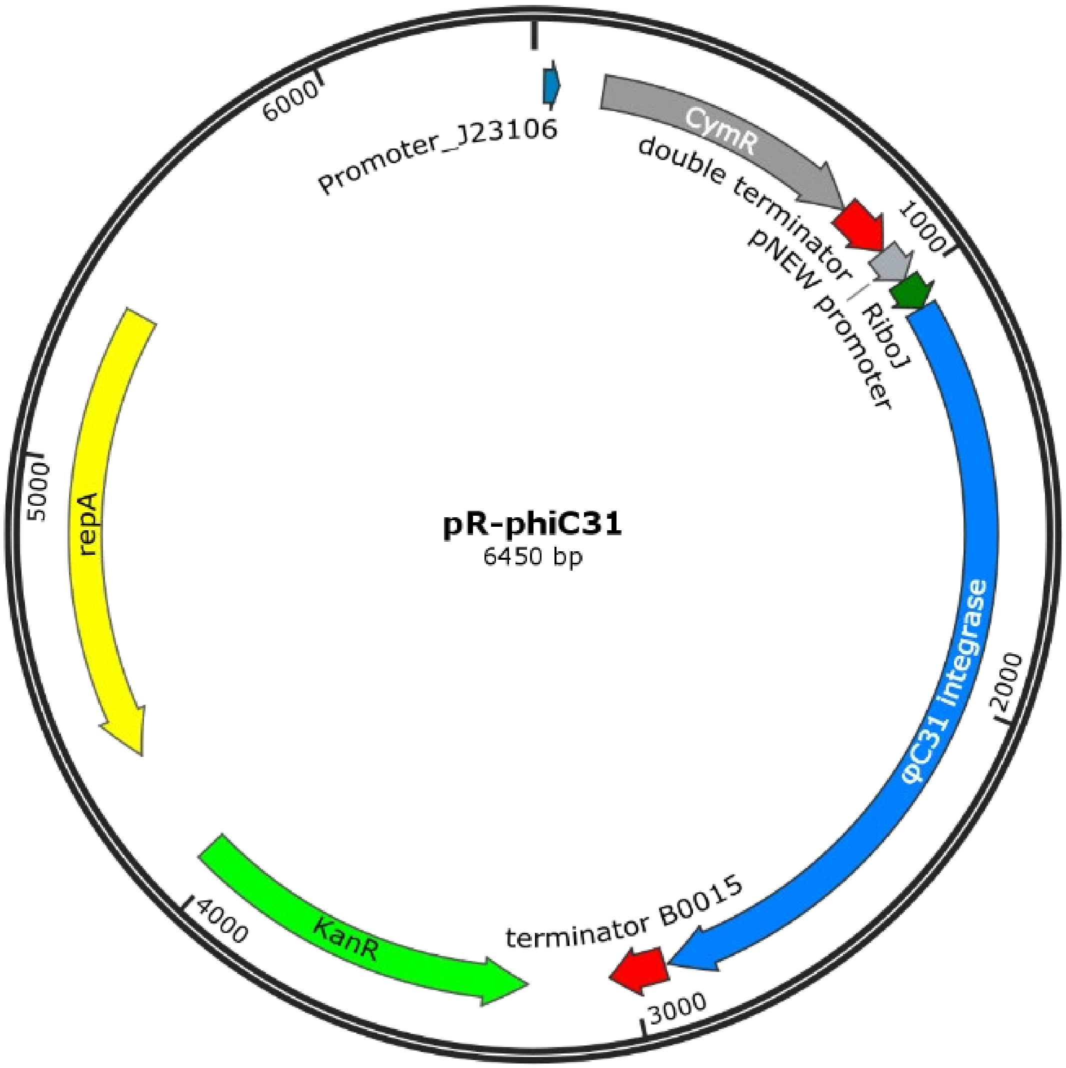

**TF Plasmids**

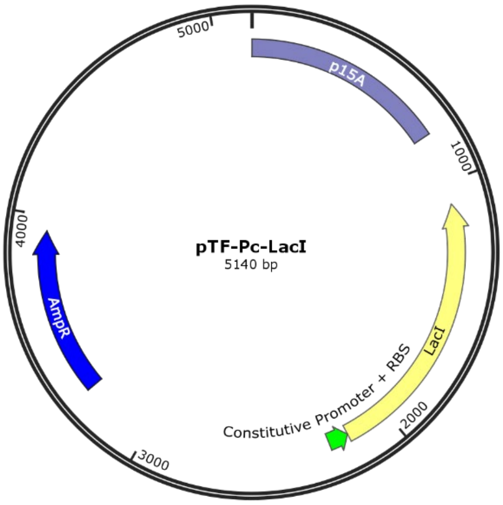
pTF-Pc-LacI
5140 bp
p15A
1000
LacI
Constitutive Promoter + RBS
2000
3000
AmpR
4000
5000

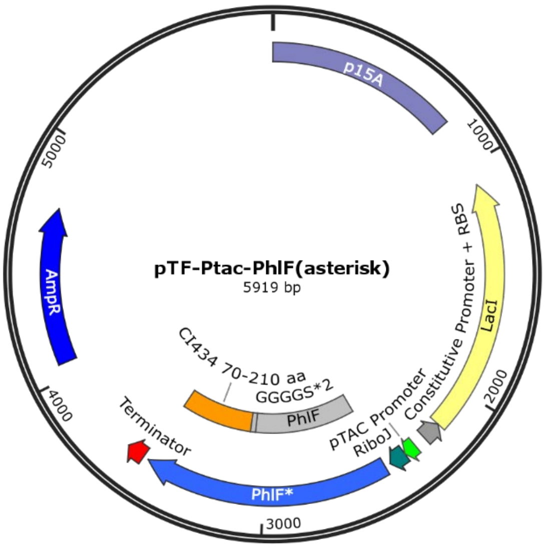
pTF-Ptac-PhlF(asterisk)
5919 bp
p15A
1000
Constitutive Promoter + RBS
LacI
2000
pTAC Promoter
RiboJ
CI434 70-210 aa
GGGGS*2
PhlF
Terminator
PhlF*
3000
4000
AmpR
5000

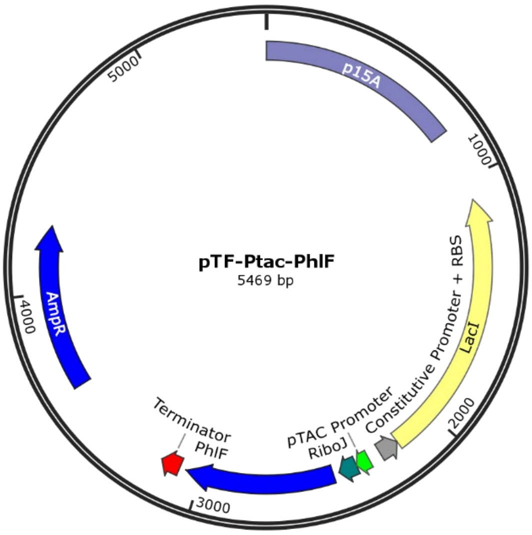
pTF-Ptac-PhlF
5469 bp
p15A
1000
Constitutive Promoter + RBS
LacI
2000
pTAC Promoter
RiboJ
Terminator
PhlF
3000
4000
AmpR
5000

**Recombinase Reporter plasmids**

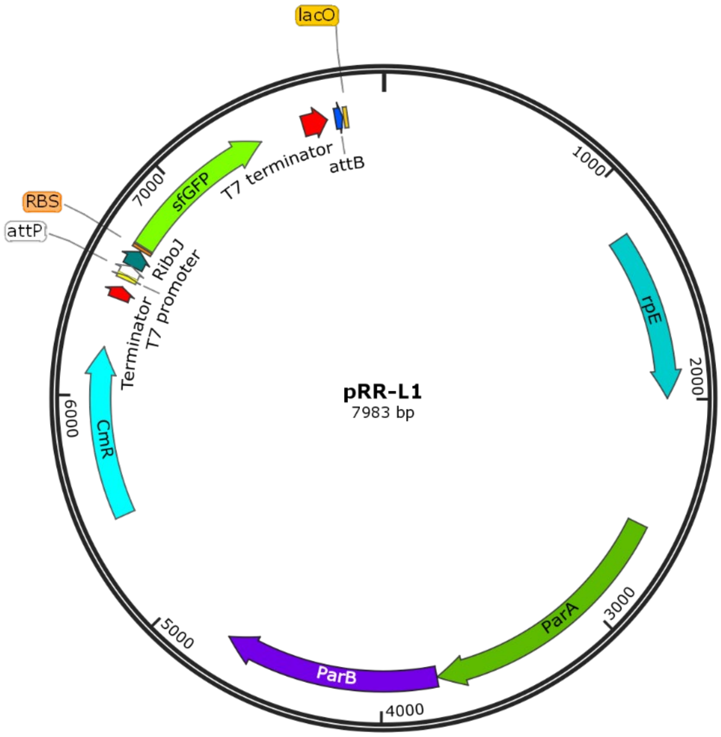


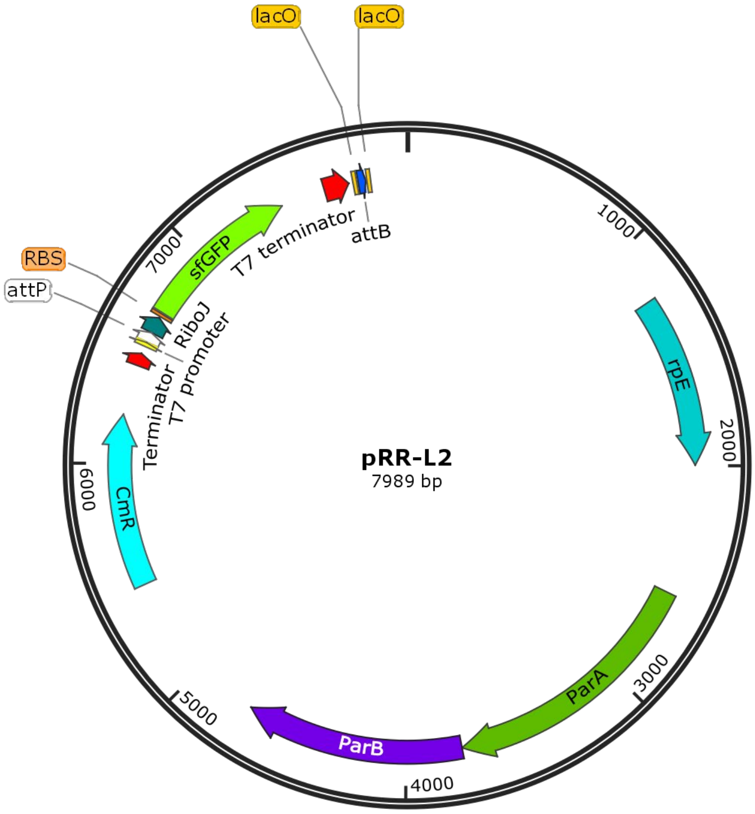

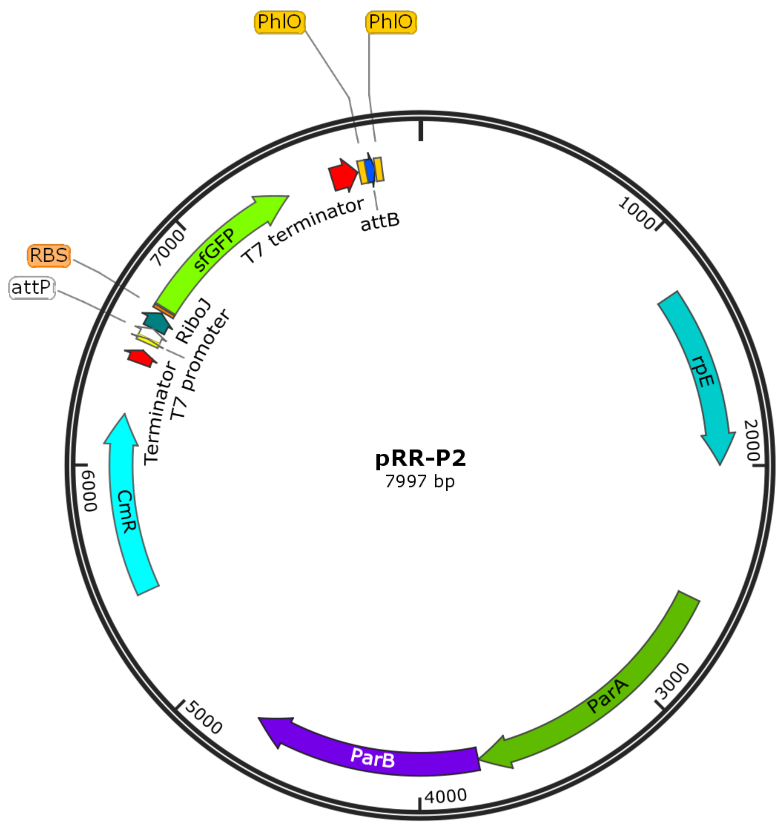


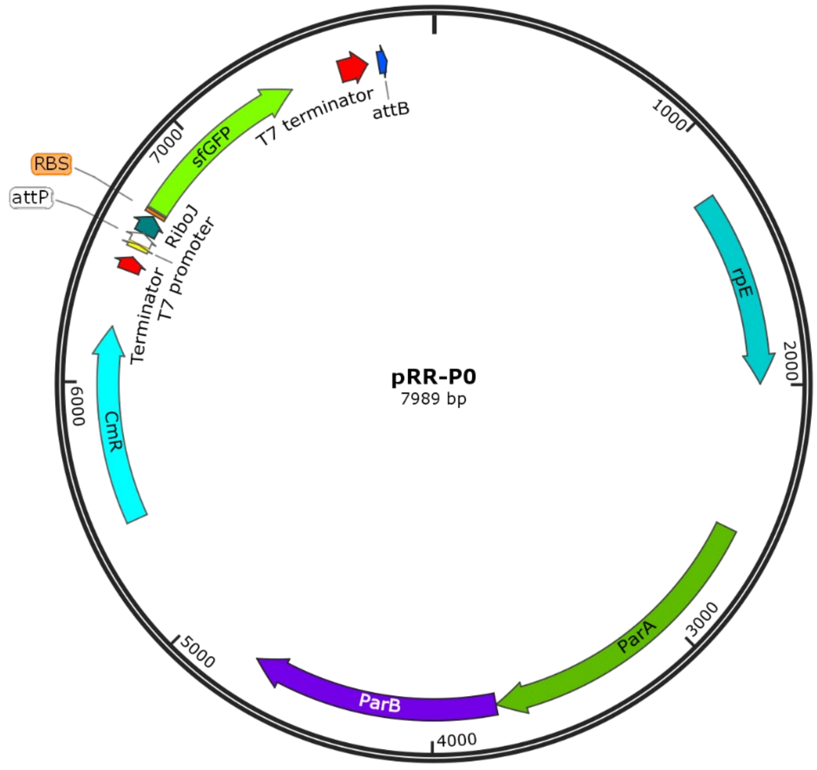


*pRR-P0 has the same sequence with pRR-L2

### Sequences of operators and attB.

| | |
|---|---|
| attB with 2 lacO operators: | ttgttatccgctcacaaggagtacgcgcccggggagcccaagggcacgccctgg<br>acccgcattgttatccgctcacaa |
| attB with 1 lacO operator: | ttctagagcaaaccgccataaaacgcccagggagtacgcgcccggggagccca[illegible]<br>gggcacgccctggcacccgcattgttatccgctcacaa |
| attB with 2 PhlO opeartors: | ATGATACGAAACGTACCGTATCGTTAAGGTcgcgccc<br>ggggagcccaagggcacgccctggcacccgcaATGATACGAAACG<br>TACCGTATCGTTAAGGT |

**Legend**: LacO attB PhlO

### The plasmids used in each experiment in this study.

| Experiment | Recombinase Plasmid | TF Plasmid | Reporter Plasmid |
|---|---|---|---|
| Proof-of-concept | pR-phiC31 | pTF-Pc-LacI | pRR-L1<br>pRR-L2 |
| Necessity of PhlF* | | pTF-Ptac-PhlF* | pRR-P2 |
| Necessity of PhlO | | pTF- Ptac-PhlF* | pRR-P0<br>pRR-P2 |
| Effect [illegible] oligomerization domain | | pTF-Ptac-PhlF*<br>pTF-Ptac-PhlF | pRR-P2 |
